# Saturating Single-Cell atlas Datasets

**DOI:** 10.1101/218370

**Authors:** Aparna Bhaduri, Tomasz J. Nowakowski, Alex A. Pollen, Arnold R. Kriegstein

## Abstract

High throughput methods for profiling the transcriptomes of single cells have recently emerged as transformative approaches for large-scale population surveys of cellular diversity in heterogeneous primary tissues. Efficient generation of such an atlas will depend on sufficient sampling of the diverse cell types while remaining cost-effective to enable a comprehensive examination of organs, developmental stages, and individuals. To examine the relationship between cell number and transcriptional heterogeneity in the context of unbiased cell type classification, we explicitly explored the population structure of a publically available 1.3 million cell dataset from the E18.5 mouse brain. We propose a computational framework for inferring the saturation point of cluster discovery in a single cell mRNA-seq experiment, centered around cluster preservation in downsampled datasets. In addition, we introduce a “complexity index”, which characterizes the heterogeneity of cells in a given dataset. Using Cajal-Retzius cells as an example of a limited complexity dataset, we explored whether biological distinctions relate to technical clustering. Surprisingly, we found that clustering distinctions carrying biologically interpretable meaning are achieved with far fewer cells (20,000). Together, these findings suggest that most of the biologically interpretable insights from the 1.3 million cells can be recapitulated by analyzing 50,000 randomly selected cells, indicating that instead of profiling few individuals at high “cellular coverage”, the much anticipated cell atlasing studies may instead benefit from profiling more individuals, or many time points at lower cellular coverage.

Recent efforts seek to create a comprehensive cell atlas of the human body^1,2^ Current technology, however, makes it precipitously expensive to perform analysis of every cell. Therefore, designing effective sampling strategies be critical to generate a working atlas in an efficient, cost-effective, and streamlined manner. The advent of single cell and single nucleus mRNA sequencing (RNAseq) in droplet format^3,4^ now enables large scale sampling of cells from any tissue, and a recently released publicly available dataset of 1.3 million single cells from the E18.5 mouse brain generated with the 10X Chromium^5^ provides an opportunity to explore the relationship between population structure and the number of sampled cells necessary to reveal the underlying diversity of cell types. Here, we present a framework for how researchers can evaluate whether a dataset has reached saturation, and we estimate how many cells would be required to generate an atlas of the sample analyzed here. This framework can be applied to any organ or cell type specific atlas for any organism.

## Results

10X Genomics recently generated an open access dataset of 1.3M cells captured and sequenced from an E18.5 mouse brain^6^ After performing quality control on the full dataset, we created randomized data subsets starting at 100,000 cells and subsampled by a factor of two down to a smallest data size of approximately 6,000 cells (SFig 1a). Each subset was randomly selected from the libraries represented in the original dataset (SFig 1b), and Louvain-Jaccard clustering generated clusters that each represented many of these libraries^7^ (SFig 1c,d). Visualization of each of these clusters in the space of the original dataset recapitulated the structure of the 1.3M cells (Fig 1a) and the proportional number of cells derived from each cluster verified that cells from all clusters were represented in every subset (SFig 2a).

**Figure 1.**
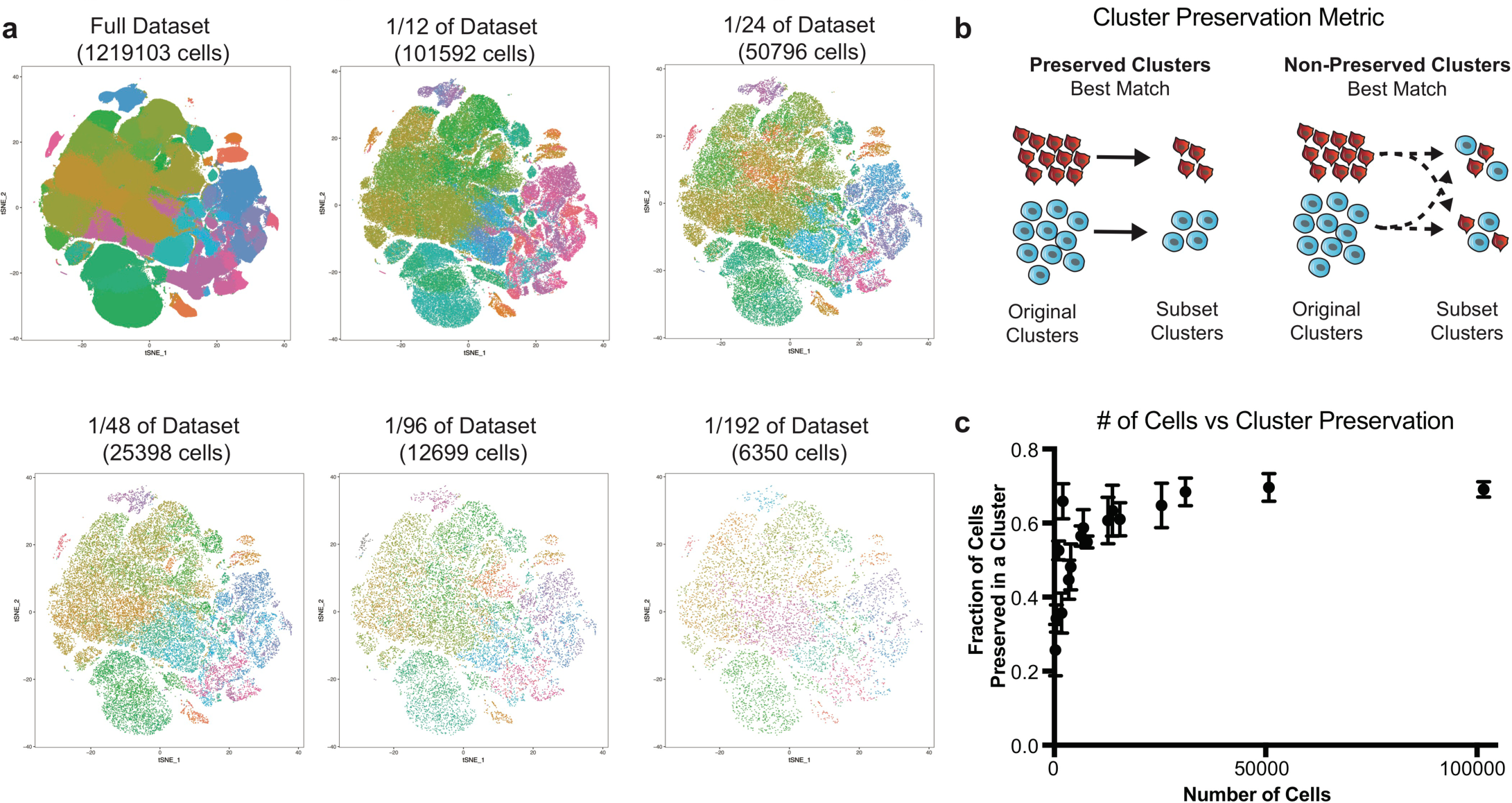
Downsampling of cell number preserves major cell type distinctions. (**a**) t-SNE plots of the full dataset and five smaller downsampled subsets. Each dataset is shown in the t-SNE space of the full dataset.Clustering was performed independently in every subset. (**b**) Cluster preservation is a key metric to evaluate similarities and differences between clusters from different analyses, measuring preservation as a fraction of the original cluster that remains in analyzed subsets. The diagram depicts a simplified cluster preservation calculation (see also Online Methods). (**c**) Cluster preservation as a function of cell number. Points are averaged within a sample from 56 downsampled subsets. The graph begins to plateau at a cell number of ~25,000 cells. For clarity, similar cell number subset preservations were averaged and standard deviation error bars added; non-averaged graph is in SFig 2B.

To compare clusters from the downsampled subsets to the clusters in the original dataset, we devised a cluster preservation metric. This metric examines how cells from the original clusters are distributed in the re-clustered subsets; the highest fraction of similarity defines the level of cluster preservation (Fig 1b). In order to explore how much cluster preservation is achieved with subsets, we scored the cluster preservation for subsets containing variable numbers of cells and observed a plateau at around 0.70, emerging around 25,000 cells (Fig 1c, SFig 2b). Interestingly, this same number was the ceiling for cluster preservation regardless of reference subset, suggesting that 30% of cells are not systematically assigned to the same cluster. These in-between cells have been previously observed in other datasets^8^ Furthermore, these findings suggest that additional cells are not useful towards recapitulating original clusters, although these findings cannot inform the “accuracy” of either clustering solution.

Different tissues or organisms may be more or less homogenous in terms of population structure, and understanding how sampling requirements differ depending on the complexity of a tissue is essential to designing effective sampling strategies. We developed a complexity index calculation that enables us to evaluate how many different cell types likely exist in a dataset by calculating Euclidean distance between cluster centroids in principal component space (Fig 2a). Because the brain is thought to be an organ comprised of particularly diverse cell types, this dataset affords a unique opportunity to explore the impact of cell population complexity on clustering and classification. We selected groups from a hierarchical tree of clusters from one of the 101,592 cell sets, intentionally generating higher and lower complexity subsets of varied cell numbers (Fig 2b, SFig 2c). While this complexity index is positively correlated to cell number, we did generate examples of lower complexity datasets with more cells (SFig 2d).

**Figure 2.**
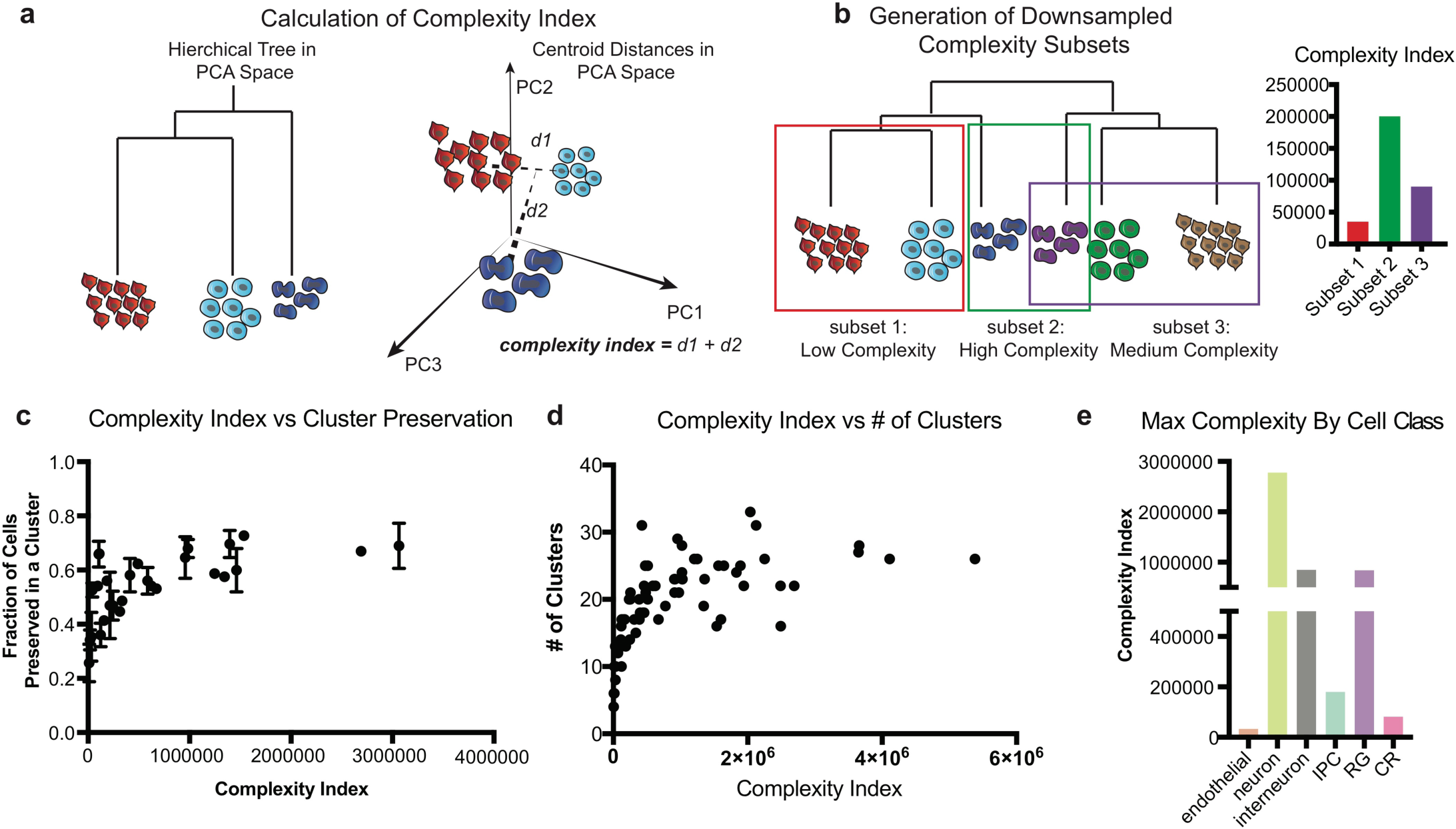
Downsampling of cell complexity preserves major cell type distinctions. (**a**) Cell complexity is calculated in PCA space of the largest reference cell set analyzed. A hierarchical tree of clusters is calculated for each subset in PCA space, and the total distance between the branches defines the cell complexity (see also Online Methods). (**b**) Cell complexity downsampling was performed by selecting branches of a larger tree with varied cell numbers and distances between groups. (**c**) Cell preservation as a function of cell complexity. Points are averaged within a sample from 56 downsampled subsets. The graph begins to plateau at a cell complexity of ~100,000. For clarity, similar cell number subset preservations were averaged and standard deviation error bars added; non-averaged graph is in SFig 2E. (**d**) Number of clusters derived from subset analyses as a function of cell complexity. The graph begins to plateau at a cell complexity of ~100,000, suggesting there is a maximal number of clusters that can be derived from a sample even as cell number and complexity increases. (**e**) Complexity calculated by cell class annotations show neurons are the most complex of the cell types retrieved.

We utilized the complexity index as an alterative to cell number, examining cluster preservation as a function of complexity. Similar to cell number, we find that the cluster preservation score plateaus at approximately 0.7, and above the complexity score of approximately 100,000, cluster preservation score did not increase (Fig 2c, SFig 2e). Interestingly, when mapping the number of clusters that are generated from an initial clustering analysis, the number of clusters similarly plateaus at the same complexity index (Fig 2d,e). These data suggest that beyond a complexity of 100,000, limited additional information about the sample can be gained through clustering analysis.

While cell preservation scores highlight how well original clusters are recapitulated, this assumes the original clusters are an accurate breakdown of the cells being analyzed. Alternatively, we devised a cell “cluster conservation” score that takes a bottom up approach, examining how the subset clusters are represented in the original dataset (Fig 3a). In general, cell cluster conservation scores are much more stable and increase only incrementally with cell number and cell complexity (Fig 3b,c). Cluster conservation can be high either when a larger original cluster gets split into multiple clusters in a subset, or when original clusters are lumped into a single cluster in the subset, suggesting a broader utility for cluster conservation in identifying biologically meaningful cell classes(SFig3).

**Figure 3.**
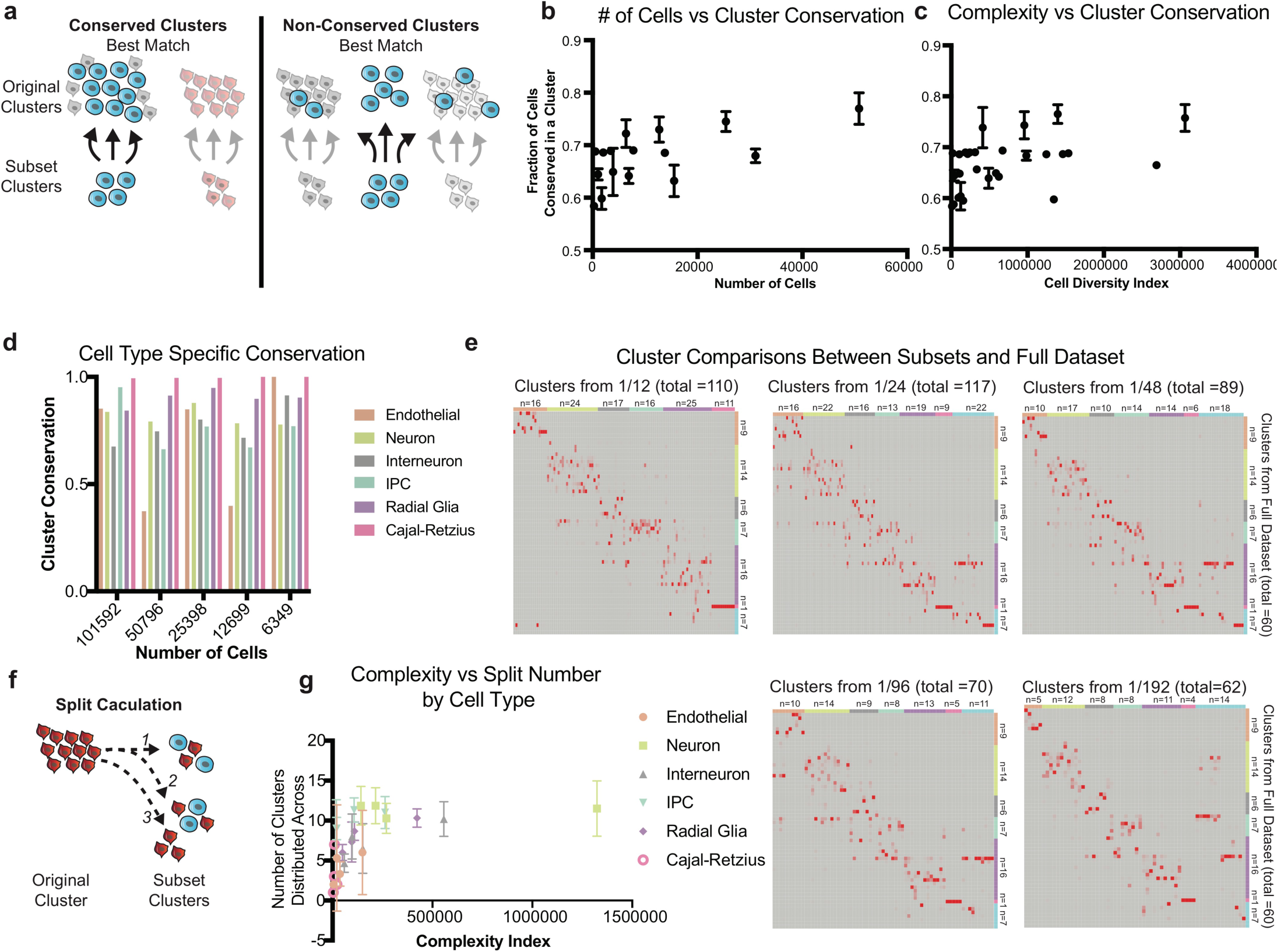
Cluster conservation from downsampled datasets. **a**) Cluster conservation is an alternative metric to evaluate similarities and differences between clusters from different analyses, measuring conservation as a fraction of the subset cluster that originates from the same cluster. The diagram depicts a simplified cluster conservation calculation (see also Online Methods). (**b**) Cluster conservation as a function of cell number. Points are averaged within a sample from 56 downsampled subsets. (**c**) Cluster conservation as a function of complexity index. Points are averaged within a sample from 56 downsampled subsets. (**d**) When grouping clusters by cell type, cluster conservation is nearly perfect for most cell types. (**e**) Heatmaps depicting the cluster conservation scores between each cell number subset and the full dataset. Blocks of conservation can be observed within each cell type, but iterative clustering results in a single cluster being split across multiple similar clusters. Interestingly, only one cluster of Cajal-Retzius cells was recovered from the full dataset, but analysis of each subset suggested that this population could be further divided into multiple clusters (see also Figure 4). (**f**) The split of single cluster can be measured by counting the number of clusters that share >= 1 cell with either the original or subset cluster, as depicted in the diagram. (**g**) Cluster split number of subset clusters as a function of complexity index divided by cell type. Again, a plateau can be seen regardless of cell type around ~100,000. More complex cell types are split more, but complexity rather than cell type appears to indicate the number of splits that may occur.

Broadly defined cell type designations are almost entirely conserved between downsampled sets and the original clustering solution (Fig 3d). Exploration of the significant PCs in each of these subsampled sets indicated that for each of the 100,000 cell sets, PCs are highly correlated, suggesting that the major sources of variation are conserved even down to the smallest subset (SFig 4). The high conservation of key sources of variation, including thegene networks defining cell types, likely explains why broad cell type assignments in the subsampled datasets remain largely the same even when substantially fewer cells are considered. However, the number of clusters in each of these initial analyses is smaller than the total number of clusters derived from the complete dataset. To identify additional subclusters, we performed iterative re-clustering (SFig 5 a-d), which substantially increased the number of clusters from each subset in proportion to the complexity of the parent cluster (SFig 5e). Interestingly, iterative clustering highlights new sources of variation (SFig 6), and the correlation of iterative PCA to the original PCA demonstrated a much smaller PC correlation.

Next, we compared clusters inferred from iterative analysis with the original clusters from the whole dataset. Surprisingly, we find that after downsampling, cells from individual clusters are reorganized into new clusters, but groups of clusters representing broad cell types are preserved.(Fig 3e). To quantify cluster stability, we measured the extent to which cells from every cluster were split by counting how many pairwise clusters contain cells from the cluster of interest (Fig 3f). Intriguingly, this cluster split measurement plateaus at a complexity of about 100,000 regardless of cell type (Fig 3g), suggesting that maximal recall of the whole dataset is capped at a cell number of ~25,000 and a complexity index of 100,000. These observations, together with the PC variance introduced by iterative clustering strongly advocate for broad cell type classification followed by targeted enrichment and subtype characterization, especially in cases where the broad survey does not yield a large cell number of a lower frequency cell classes.

The metrics proposed here characterize the efficacy of varied downsampled subsets in recapitulating initial clusters, but none of the metrics indicate the sampling or clustering strategy most effective in recovering biologically interpretable clusters. To better understand the nature of downsampling, we focused our analysis on Cajal-Retzius (CR) cells, one of the lowest frequency cell types in the forebrain. CR cells are essential to the laminar organization of the brain^9,10^ and have been characterized to originate from several sources within the brain that impart them with appropriate transcriptional markers of origin^11^. To explore this cell type, we isolated cells in the *Reln+, Tbr1+* cluster from the full 1.2 million cells dataset. By iteratively clustering these cells, we identified 18 distinct clusters with at least 10 marker genes distinguishing each cluster (Fig 1a, SFig 7a,b). The same process was applied to CR cells from each of the downsampled subsets originating from one 100,000 cells matrix.

Analysis of the clusters resulting from whole set iterative clustering suggested that some clusters were enriched for the highest and lowest levels of mitochondrial content as a fraction per cell (SFig 7c), and some had no unique identifiers separating them from other clusters, only a combination of marker level differences (SFig 7d). Other clusters did have unique marker genes, though most genes were lost as markers through the downsampling process (SFig 7e). However, two groups of clusters did highlight *Foxg1* and *Lhx9*^12,13^, markers indicating putative developmental structure of origin. Violin plots of the expression of these genes in the full dataset and the downsampled sets show that while *Lhx9* maintains distinct cluster specific expression throughout downsampling, *Foxg1* loses cluster enrichment below 1/24^th^ of the dataset (~25,000 cells, 815 CR cells). Together, this may indicate that while a certain minimum number of cells is necessary to recover some cell type distinctions, not every cluster may be biologically relevant. Instead, technical factors may influence variation during exhaustive iterative clustering, even after stringent quality control. Nonetheless, it is possible that the 9 CR clusters from the full dataset without clear markers are biologically important. Similarly, CR clusters from subsets smaller than 1/24^th^ of the dataset may have biological meaning, but we were unable to elucidate clear, meaningful distinctions.

The CR subset offers an opportunity to explore how data subsets compare to the curves we developed with our various metrics. By plotting these CR subsets along the trajectories of cluster preservation, complexity, number of clusters, and split curves, we observe that by these metrics, the analysis has not yet reached saturation (Fig 4d). However, our biological interpretation suggests that the main cell type subsets are identifiable within the subsets analyzed here. Our analysis provides a pragmatic analytical framework for evaluating whether a single cell dataset has been saturated. Specifically, downsampling of the dataset followed by implementation of the cell preservation score and complexity index analysis can reveal whether cluster diversity is saturated. If a linear regression on a plot of number of cells or complexity versus cluster preservation fits the downsampling with an R-squared less than 0.6 and progressively decreases from larger cell number, saturation may be reached (Fig 4e-f). This analysis suggests that saturation is much more quickly reached from the perspective of broad cell types, while iterative subtype identification is more fluid and requires careful biological validation.

**Figure 4.**
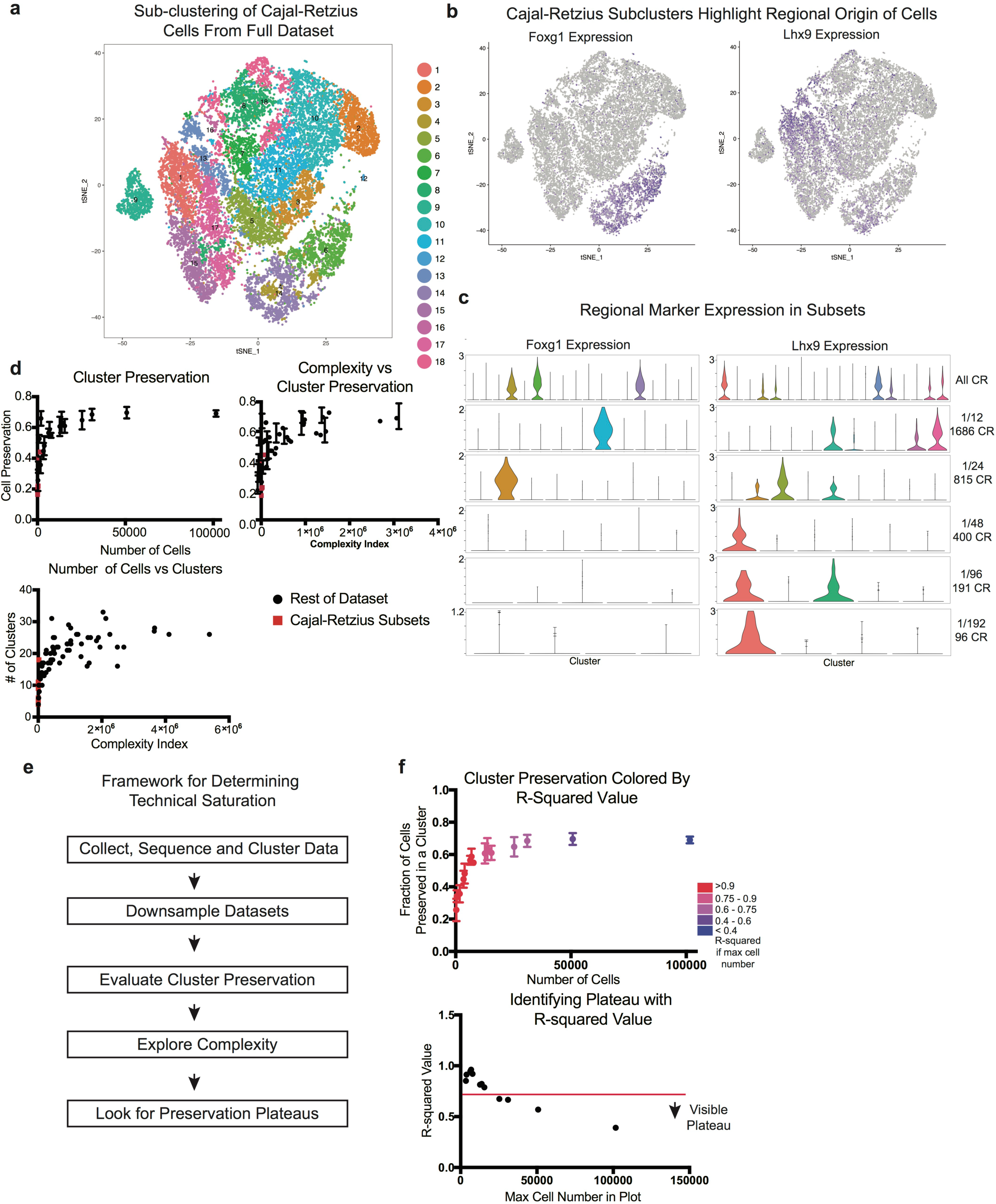
Downsampling of Cajal-Retzius Cells. (**a**) t-SNE plot depicting the iterative clustering result of all 20,550 Cajal-Retzius (CR) cells from the full dataset. (**b**) Regional origin is a well studied classifier of CR subtypes, and two of these markers feature prominently in the iteratively clustered dataset: *Foxg1*is enriched in 3 clusters while *Lhx9* is enriched in 7 clusters. (**c**) Violin plots of regional markers in the full datasets and CR subsets of downsampled datasets indicate that these markers are enriched in one more clusters up until 1/24 of the dataset is sampled, after which *Foxg1* enrichment is diluted across multiple clusters. *Lhx9* enrichment is conserved to even the smallest downsampled subset. (**d**) Enrichment metrics of CR cells in the context of previously shown metrics indicate that informatically, saturation of this cell type has not yet been achieved. (**e**) Framework to evaluate if technical saturation has been achieved. (**f**) Examination of R-squared values when incrementally decreasing the number of maximum cells used in the analysis shows that plateau emerges around an R-squared value of 0.6.

## Discussion

Here, we present a framework for evaluating if cell atlas datasets have reached saturation by introducing a few simple and practical principles to evaluate whether enough cells have been sampled to characterize a heterogeneous tissue. Preservation, a metric for how well original clusters are preserved in data subsets, plateaus around 25,000 cells in the full dataset of 1.3 million cells from the E18 mouse forebrain. The plateau does not approach 1, but PCA beyond major subtypes likely introduces technical noise into the analysis, suggesting that clustering beyond this level of preservation may be an analytical artifact. This viewpoint is bolstered by the lack of biological insight and lack of distinctmarkers in clusters derived from iterative clustering of the Cajal-Retzius subsets.

We additionally present complexity index, which scores the relative heterogeneity of a sampled cell population. We note that beyond a certain complexity, cell preservation also plateaus, at around 100,000 in our analysis. This plateau suggests that clustering has a maximal number of divisions that can be generated in a dataset per analysis, and supports a role for iterative clustering. However, even with iterative clustering, cluster preservation plateaus. This indicates that only a certain number of meaningful divisions can be identified within any dataset, and that effective generation of the cell atlas will require multiple iterations of data collection, saturation analysis, and tissue level validation. In addition, the analytical framework presented here provides a model for researchers to explore their own datasets and utilize data-driven strategies to evaluate when cell number and complexity have been achieved.

## Methods

### Data Management

To access the full matrix of the 1.3M cell set, we used the python instructions for accessing the HD5 object. The CellRangersoftware that is used to process 10X data outputs three files for genes, matrix, and barcode. Because of the size limitations of the full dataset, we broke the dataset into 64 parts and wrote these subsets to file. We then used the Seurat^3^ Read10X command to assemble these three files in R. For each subset, we normalized the total counts per cell to 10,000, eliminated cells with > 10% mitochondrial or ribosomal content as well as those with fewer than 1000 genes per cell. This quality control filtration reduced the total cell number in the analysis to 1,219,103.We evenly combined these filtered matrices into 4 parts, and then into 2 parts. Using a server node with 64 cores, a 2.6 GHz processor, and 512 GB of RAM, any additional sized matrices resulted in the error “problem too large”. Therefore, matrices of approximately ~100,000 cells were the unit primarily used in this analysis.

### Downsampling

Random downsampling to the 101,592 cell sets occurred by repeated (>10 times) shuffling the two matrices of the full data, cutting them each in half, and recombining to make 2 random matrices. Once this had been performed the 101,592 cell sets were generated in order from the full dataset, without replacement. Subsequent downsampling to smaller datasets resulted from random selection of this dataset; each subset formed the full set of available cells for the next downsampling (i.e, 50,796 cells were taken from one set of 101,592 cells, and the next subset of 25,398 cells was sampled from the set of 50,796 cells). Twelve 101,591/101,592 cell sets were generated, and 9 were analyzed for the sampling parameters included in the main figures.

### Clustering

The full dataset was clustered by 10X Genomics using their CellRanger v1.2 graph based clustering solution, loupe browser, and industrial scale computational resources. Clustering was performed on 100,000 cell sets and smaller using the graph based Louvain-Jaccard method that has been previously described (Shekhar et al, 2016). Briefly, fastPCA is performed on the full centered and scaled expression matrix for 50 principal components. Using the formula laid out by Shekhar et al, we calculated the number of significant PCs to be 18 and used this number of PCs for all comparative analyses. A nearest neighbors calculation is performed in PCA space for 10 nearest neighbors, and from this analysis a Jaccard distance was calculated for each pair of neighbors. This data is used as the input to the Louvain clustering. t-SNE plots were either generated in the PCA space of each analysis, or the loupe t-SNE or other clustered plots were used as the reference coordinates for a smaller dataset.

Iterative clustering was performed in the same way, but clusters of the same type were grouped together and used as the input to the clustering process.

### Cluster Preservation

Because this analysis uses subsets of cells with the same names as the analyses the clusters are begin compared to, cluster preservation was calculated by counting the number of cells that were the same in each pairwise cluster comparison. The fraction was calculated using the number of cells in the smaller dataset’s cluster. Preservation for each original cluster, then, was the maximum of the compared fractions across all subset clusters.

### Complexity Index Calculation

Complexity index is a way of measuring relative sample complexity by using Euclidean distances in PCA space. For each set of clusters, the Seurat BuildTree function was used to hierarchically represent clusters in the PCA space of the cells being analyzed. Complexity index was calculated in the largest available PCA space of the cells being analyzed (i.e. 25,398 cells used the PCA of the corresponding 101,592 cell set). To generate the index, the branch lengths of the tree were added based upon the centroid distances in PCA space. The index is a number in arbitrary units. Complexity index scales positively with cell number, but it is possible to generate larger cell numbers with smaller complexity scores. Downsampling of complexity was performed by picking high and low complexity subsets from a tree of clusters from one of the 101,592 cell sets.

### Cluster Conservation

Cluster conservation is a “bottom up” evaluation of cluster preservation. Instead of observing how intact original clusters are in the subsets, cluster conservation measures the maximal correspondence in terms of fraction of cells the same in pairwise cluster comparisons from the perspective of the new clusters. It uses the same calculation as cluster conservation but is the reciprocal analysis. Cluster conservation can be high even with low cluster preservation, particularly when there is a large imbalance of the number of clusters in the two analyses being compared.

### Split Calculation

Measuring the number of splits is done by simply counting the number of pairwise clusters that have any cells from a cluster of interest, identifying how split up a cluster is. For example, even if cluster preservation is only 0.50, but its cluster split is 2, then it is a single cluster evenly split and could be considered a strongly preserved cluster. However, a cluster with preservation of 0.5 but a split of 10 would be considered to be much less preserved as its cells are found in a wide variety of subsequently generated clusters.

---

**Supplementary Figure 1.**
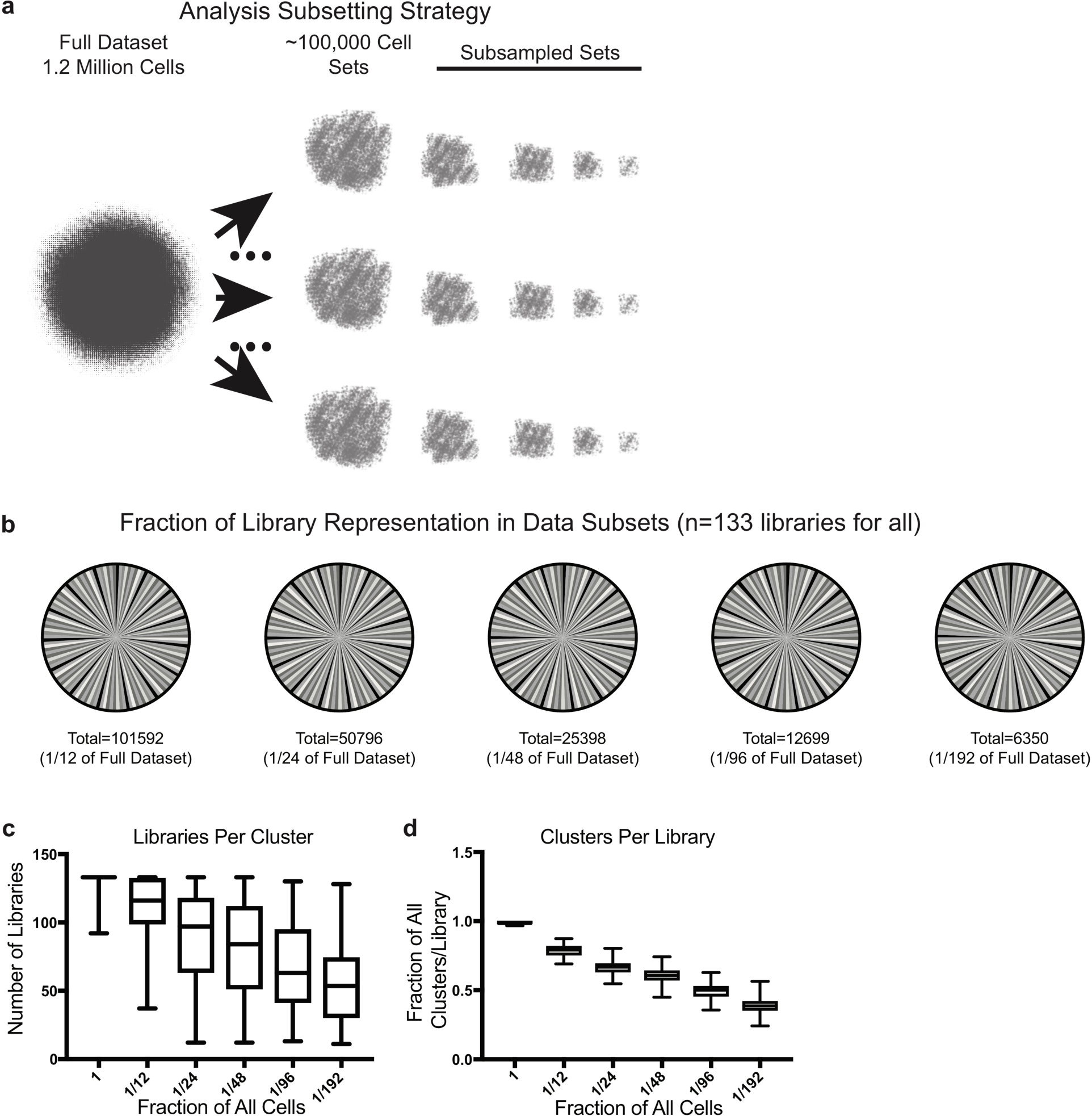
Library and cluster composition metrics (**a**) Schematic depicting subsampling strategy for analysis (**b**) Library representation in each subset used in subsequent analyses. Each of the libraries is equally represented in all datasets. (**c**) Number of libraries in a cluster across subsampled datasets. This number decreases as the dataset gets smaller, but the majority of clusters still have 10s of libraries even in the smallest set. (**d**) The fraction of clusters per library. All libraries contain a diversity of cell types, and much of this representation is preserved with downsampling.

**Supplementary Figure 2.**
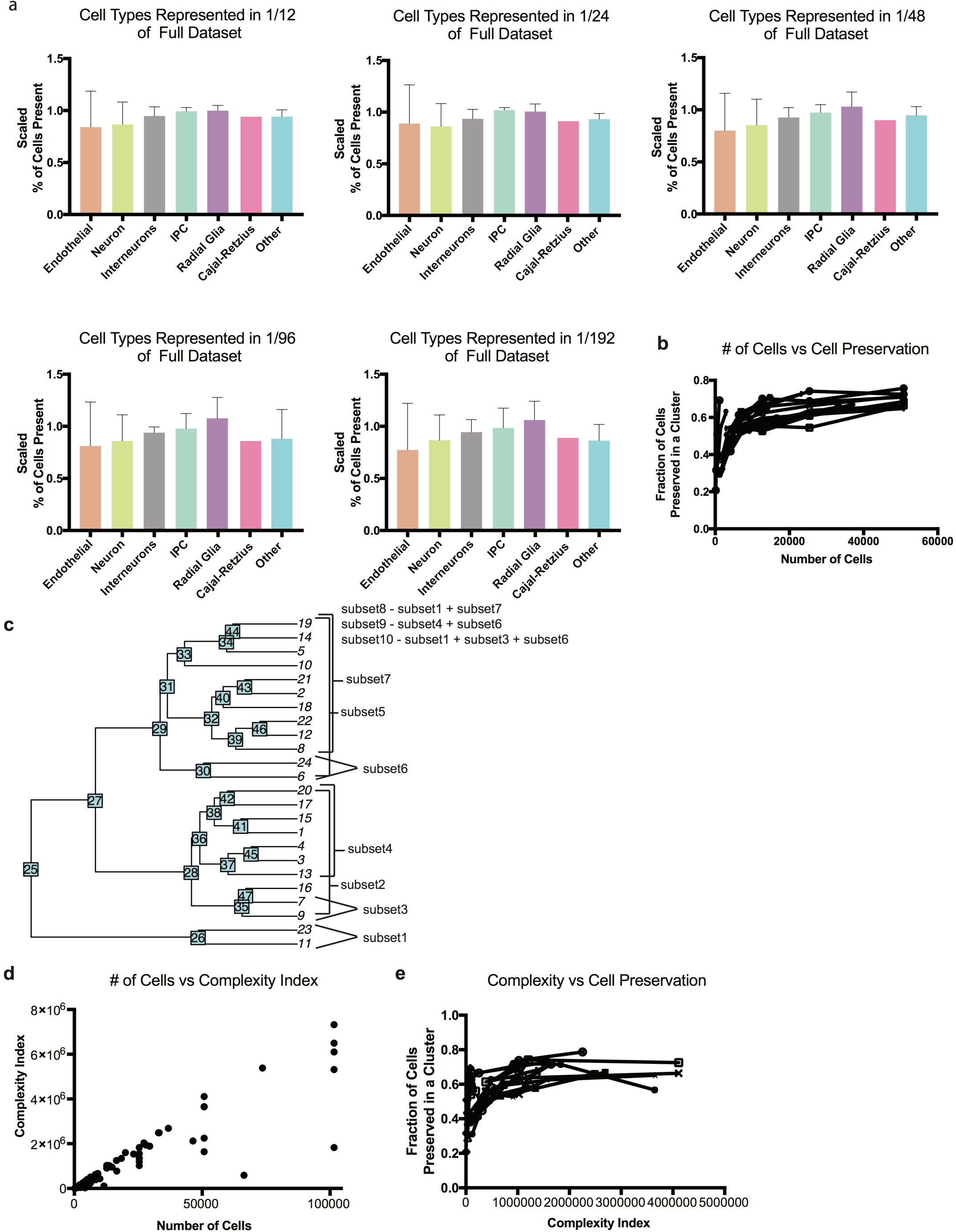
Complexity index scales with downsampling. (**a**) Scaled proportion of cell types represented in each of the downsampled datasets. For the complete dataset, cell types were annotated by examining cluster markers and assigning cell type where possible. The number of cells from each of the full dataset represented in each subset was use to generate a fraction, and scaled to the proportion of the full dataset present in each subset. (**b**) Cluster preservation scores are represented in Fig 1C, but showing lines to indicate downsampled datasets. (**c**) A hierarchical tree of clusters from one set of 101592 cells. This hierarchy was used to generate subsets of intentionally varied cell numbers and complexities. (**d**) Plot of number of cells versus complexity index. In general, cell number is correlated to complexity but complexity can be less in a larger number of cells, particularly when downsampling. (**e**) Plot of complexity versus cell preservation, as shown in Fig 2C, but with lines to indicate downsampled samples from the same set.

**Supplementary Figure 3.**
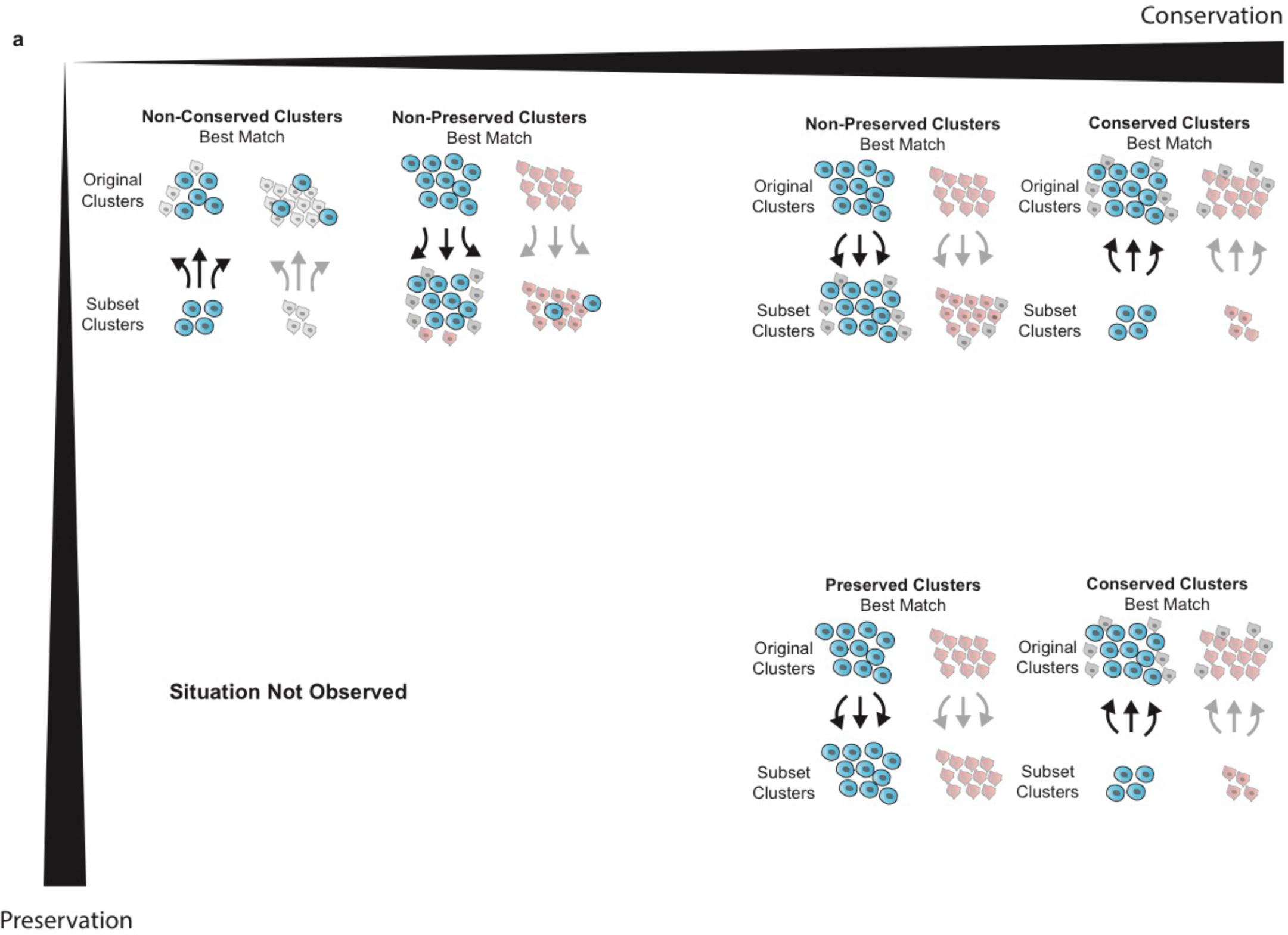
Schematic of conservation versus preservation metrics. (**a**) Preservation and conservation are two concepts that can be used to measure cluster fidelity between datasets. Preservation is more dynamic and can range from low to high, but lower conservation is also possible. Schematic depicts the nature of the three intersections of preservation versus conservation that are observed in this dataset.

**Supplementary Figure 4.**
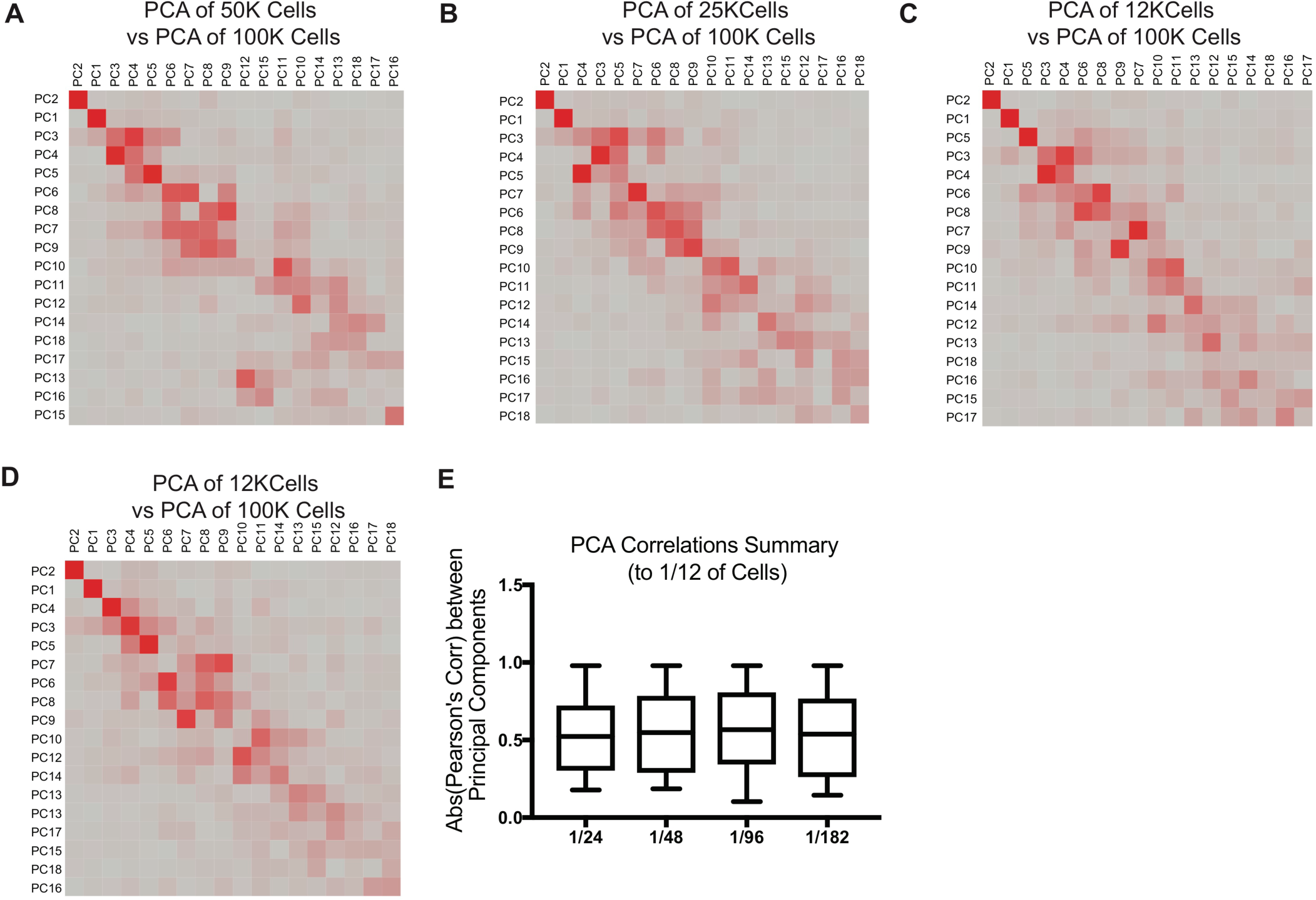
Major sources of variation are preserved with downsampling. (**a-d**) When comparing the 100K cell matrix to further downsampled subsets, a strong diagonal across the matrix is observed. The PCA conservation, though not one-to-one, indicates strong principal component preservation across datsets. (**e**) Quantitative summary of the absolute values of the best PC correlations between the datasets explored here.

**Supplementary Figure 5.**
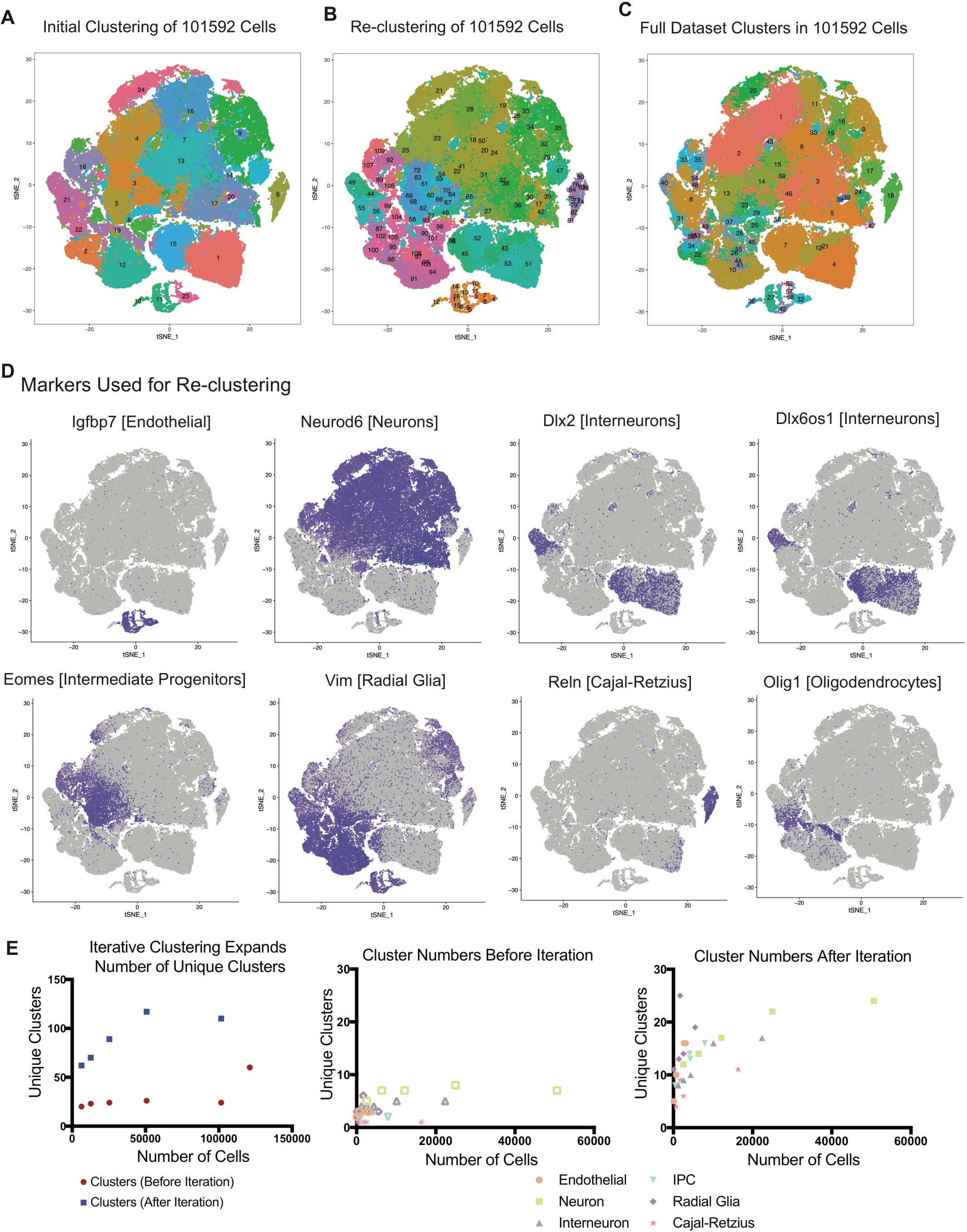
Major cluster features and subgroup determination. (**a**) tSNE plot showing the initial clustering of one dataset of 101592 cells. (**b**) Using iterative reclustering, tSNE plot is colored by new clustering analysis. (**c**) Using the cluster designations of the cells used in this subset, clusters are colored on tSNE plot by their loupe cluster annotations. (**d**) Cluster markers used to designate clusters for iterative clustering analyses. (**e**) Plots of number of clusters identified before and after iterative clustering analyses. The number increases to a point, but the maximum is actually seen at 50K cells. Segregating this representation by cell type indicates that additional cluster resolution is dependent upon the number of cells in the subtype being re-clustered.

**Supplementary Figure 6.**
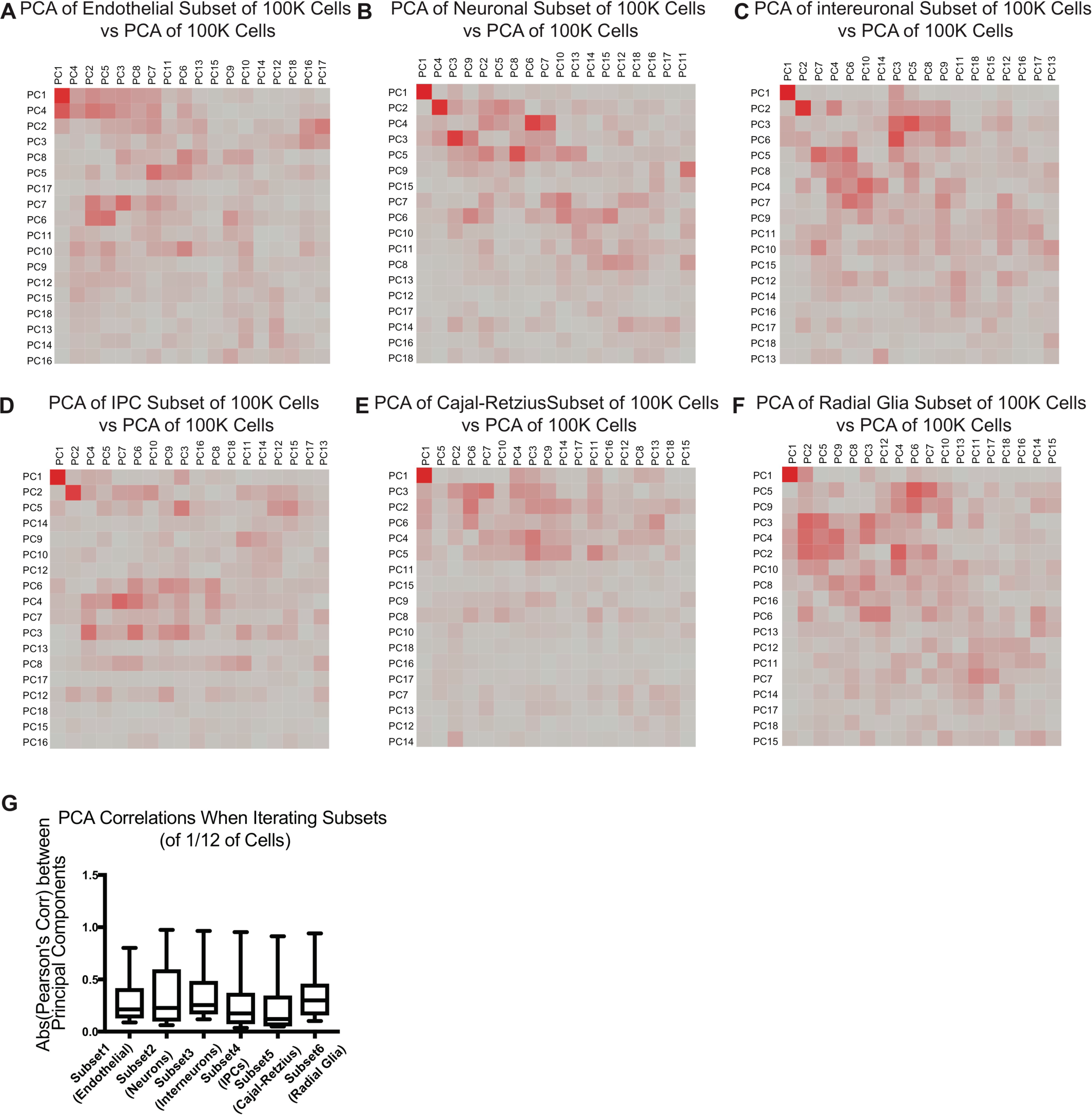
New sources of variation emerge in data subsets. (**a-f**) Examination of PCA correlations between the whole dataset of 101592 and the cell type specific subsets being analyzed shows that only one or two PCs are typically highly correlated to one another, indicated the iterative clustering introduces additional sources of variation. (**g**) Quantitative summary of the of the absolute values of the best PC correlations between the datasets explored here.

**Supplementary Figure 7.**
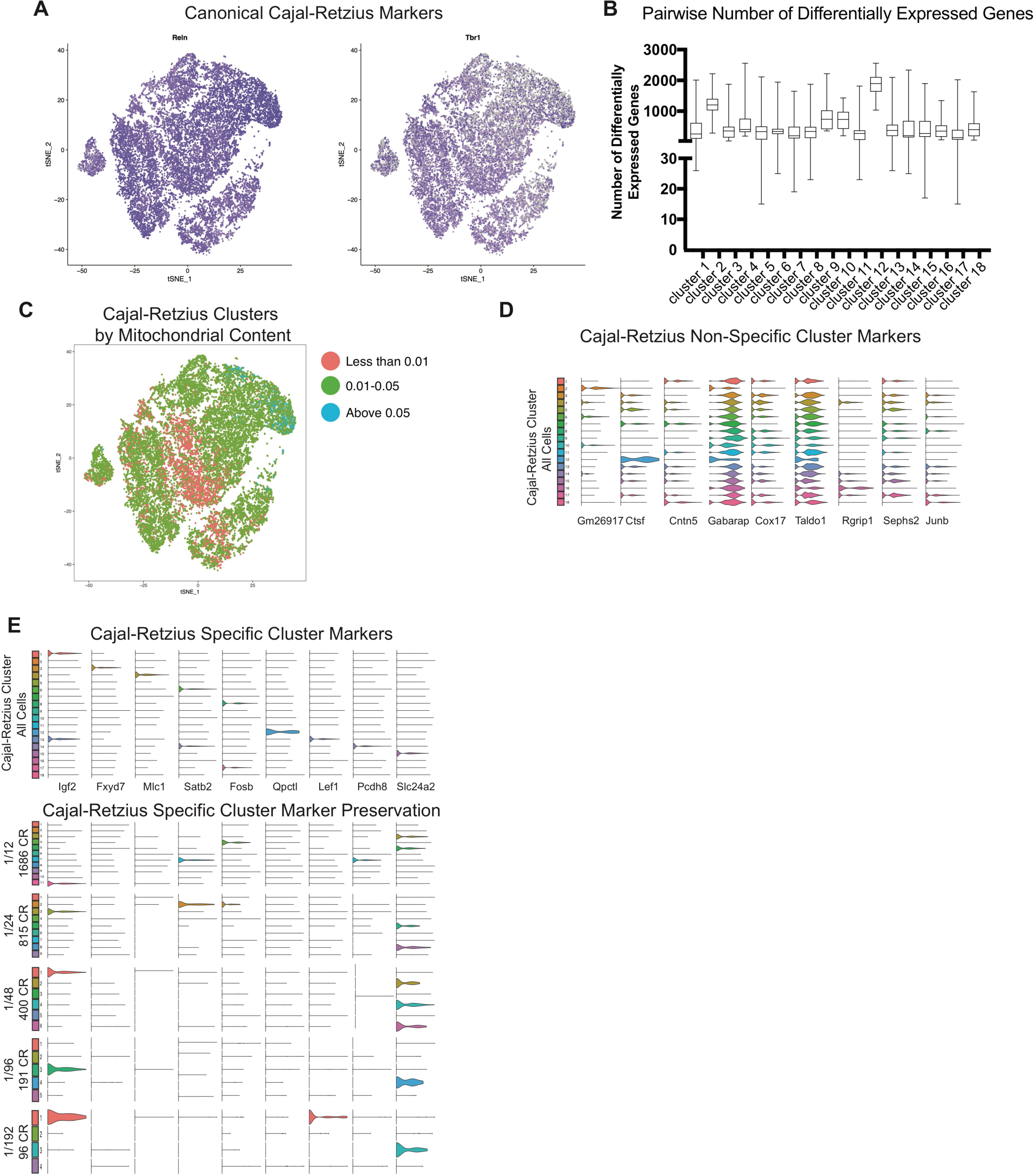
Cajal-Retzius cell diversity. tSNE plot of 20K CR cells from whole dataset colored by *Reln* and *Tbr1* expression indicating the clustering isolated canonically marked CR cells. (**b**) Box plots of the number of differentially expressed genes between each cluster and all other clusters indicates that the 18 clusters identified are informaticaly distinct. (**c**) Coloring CR tSNE plot indicates pockets of enrichment of clusters dependent upon mitochondrial content, even after QC filtering. (**d**) Violin plots of non-specific CR cluster markers, these markers were the best enriched for some clusters but show expression across multiple clusters. (**e**) Specific CR cluster markers are shown in the top half, these markers are strongly enriched in one or clusters of the iteratively cluster 20K CR cells from the whole dataset. Examination of marker expression of these genes in downsampled CR sets shows *Igf2, Satb2, Lef1, and Slc24a2* are largely conserved with downsampling but other markers are lost, sometimes with the first downsampled set.

